# MinION-in-ARMS: Nanopore Sequencing To Expedite Barcoding Of Specimen-Rich Macrofaunal Samples From Autonomous Reef Monitoring Structures

**DOI:** 10.1101/2020.03.30.009654

**Authors:** Jia Jin Marc Chang, Yin Cheong Aden Ip, Andrew G. Bauman, Danwei Huang

## Abstract

Autonomous Reef Monitoring Structure (ARMS) are standardised devices for sampling biodiversity in complex marine benthic habitats such as coral reefs. When coupled with DNA sequencing, these devices greatly expand our ability to document marine biodiversity. Unfortunately, the existing workflow for processing macrofaunal samples (>2-mm) in the ARMS pipeline—which involves Sanger sequencing—is expensive, laborious, and thus prohibitive for ARMS researchers. Here, we propose a faster, more cost-effective alternative by demonstrating a successful application of the MinION-based barcoding approach on the >2mm-size fraction of ARMS samples. All data were available within 3.5–4 h, and sequencing costs relatively low at approximately US$3 per MinION barcode. We sequenced the 313-bp fragment of the cytochrome c oxidase subunit I (COI) for 725 samples on both MinION and Illumina platforms, and retrieved 507–584 overlapping barcodes. MinION barcodes were highly accurate (~99.9%) when compared with Illumina reference barcodes. Molecular operational taxonomic units inferred between MinION and Illumina barcodes were consistently stable, and match ratios demonstrated highly congruent clustering patterns (≥0.96). Our method would make ARMS more accessible to researchers, and greatly expedite the processing of macrofaunal samples; it can also be easily applied to other small-to-moderate DNA barcoding projects (<10,000 specimens) for rapid species identification and discovery.

## 1. Introduction

An estimated 80% of living species remain unknown to science, including up to 90% of the world’s marine species (Mora et al., 2011; Appeltans et al., 2012; Wilson, 2017). In the context of accelerating global change, there is an urgent need to more rapidly discover and assess species diversity, given that rates of species losses are predicted to occur faster than we can document them (Costello and Wilson, 2011). Fostered by the need to expedite species discovery, marine researchers have proposed the use of the Autonomous Reef Monitoring Structure (ARMS), a standardized sampling tool that enables comprehensive documentation of marine biodiversity beyond standard indicator species (Plaisance et al., 2011a; Leray and Knowlton, 2015). Briefly, ARMS are designed to mimic the structural complexity of coral reefs and are commonly deployed on the marine benthos for a length of time to allow marine organisms to colonise before subsequent retrieval (Knowlton et al., 2010). All organisms on the units are then subjected to DNA barcoding for species identification and quantification (Leray and Knowlton, 2015). Globally, approximately 1,700 ARMS units have been deployed under the Global ARMS Program (https://www.oceanarms.org/getting-involved), an initiative helmed by the Smithsonian National Museum of Natural History that encourages the deployment of ARMS around the world with the aim of consolidating and making all ARMS-related data available. Notably, ARMS have become a widely-utilised method for assessing benthic diversity in many shallow marine systems (Plaisance et al., 2011a, 2011b; Leray and Knowlton, 2015; Al-Rshaidat et al., 2016; Hurley et al., 2016; Pearman et al., 2016, 2018, 2019; Pennesi and Danovaro, 2017; Ransome et al., 2017; Carvalho et al., 2019; David et al., 2019; Hazeri et al., 2019).

One of the most common criticisms of utilising ARMS is the high sequencing costs (Danovaro et al., 2016). A standard ARMS sequencing workflow (see Leray and Knowlton, 2015) involves removing all fauna from ARMS and sorting into either sessile or motile fractions, with the motile fraction being further subdivided into 3 distinct size-ranges (i.e. >2-mm, 500-µm–2-mm, 106–500-µm; Figure 1). Specimens sorted into the largest size range (>2-mm) of the motile fraction are barcoded via Sanger sequencing while the other fractions undergo metabarcoding (Figure 1). While the former is done to retain sample-sequence association for the >2-mm size fraction (Leray and Knowlton, 2015), it has never been clear why Sanger sequencing was chosen over more advanced barcoding methods (i.e. next-generation sequencing (NGS); Shokralla et al., 2014; Meier et al., 2016), especially given that other size fractions already undergo NGS metabarcoding (Leray and Knowlton, 2015). Additionally, NGS has already proven to have near-perfect accuracy (≥99%) when benchmarked against Sanger sequencing (Baudhuin et al., 2015; Beck et al., 2016). Sequencing, however, typically requires access to a well-equipped molecular laboratory (Glenn, 2011; Quail et al., 2012), and Sanger costs remain high at US$18 per sample (Meier et al., 2016), making ARMS sequencing an expensive endeavour. Consequently, researchers have resorted to using imaging techniques and morphological examination of the ARMS plates as an alternative to assessing biodiversity (Hurley et al., 2016; David et al., 2019), despite morphology-based approaches being less cost-effective (Hayes et al., 2005). Sanger sequencing costs may also be the reason there is a decreasing representation of >2-mm size fractions (Supplementary Table 1). Fortunately, sequencing technologies are rapidly improving, and the rise of third-generation sequencers aimed at democratizing sequencing to the masses (Mikheyev and Tin, 2014) are now challenging the dominance of second-generation mainstays like Illumina. One such innovation is the MinION sequencer.

**Figure 1.**
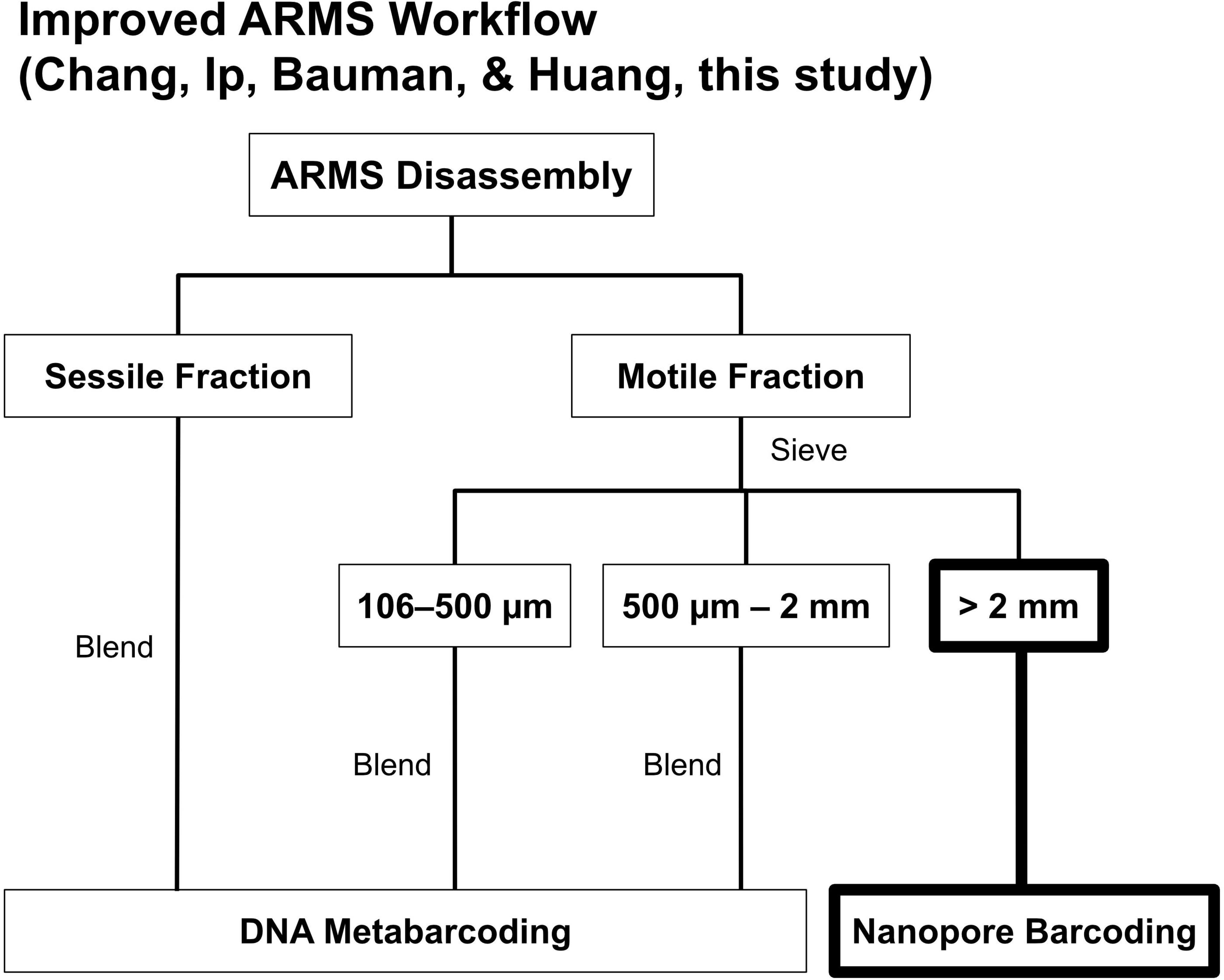
The ARMS processing workflow by Leray and Knowlton (2005). We replaced Sanger sequencing of >2-mm specimens with nanopore barcoding highlighted in **bold**.

The MinION is a small handheld sequencer that was introduced in 2014 by Oxford Nanopore Technologies (ONT). MinION’s release was significant for nucleic acid sequencing for several reasons: (1) its lower entry and per base sequencing cost, compared to second-generation sequencing technologies, (2) its ability to perform long-read sequencing, which is ideal for genome assemblies, (3) its compact size and portability, and (4) its ability to generate data real-time (Mikheyev and Tin, 2014). However, despite these promising advantages, nanopore sequencing remains hampered by its fairly high raw read error rate. For example, raw MinION read accuracies were 65–88% in its initial launch phase (Lu et al., 2016), but ongoing improvements in flow cell chemistry have increased the average read accuracy to ~90% for the R9.4 flow cells (Tyler et al., 2018; Wick et al., 2019). Further, new error-correction pipelines for barcoding have since emerged, including the *miniBarcoder* (Srivathsan et al., 2018, 2019), *minibar* (Krehenwinkel et al., 2019), and ONTrack (Maestri et al., 2019), which allow users to capitalize on nanopore sequencing advantages while keeping error rates low.

Given the recent advances in sequencing technology, we propose that the MinION-based barcoding approach can also be applied to ARMS research, specifically to process >2-mm samples in place of Sanger sequencing. We demonstrate this through application of the *miniBarcoder* pipeline (Srivathsan et al., 2018) on >2-mm samples collected from ARMS units deployed in Singapore. This pipeline was chosen because its utility has been thoroughly demonstrated on insects (Srivathsan et al., 2018, 2019) and seafood (Ho et al., 2020), and here we extend its applicability to a more taxonomically diverse sample set. We posit that the MinION method is more cost-efficient and will enable a much faster ‘*sample-to-sequence*’ workflow for ARMS researchers, with the added advantage of scalability depending on individual project requirements, making it a highly useful pipeline for any small-to-moderate (<10,000 samples) specimen-rich biodiversity barcoding endeavours.

## 2. Materials and Methods

### 2.1. Deployment, retrieval and processing of ARMS units

Four sets of three ARMS units were deployed across four sites in the Southern Islands of Singapore for two years from July 2016 to July 2018. The ARMS units were processed according to Leray and Knowlton (2015). Collections were authorised by the National Parks Board (permit number NP/RP15-088). The >2-mm samples were vouchered individually and classified by morpho-phylum during the disassembly and sorting phase. Classifications were recorded for downstream congruence checks with molecular data (see below). Samples were handled according to NUS Institutional Animal Care and Use Committee (IACUC) guidelines (IACUC Protocol B15-1403).

### 2.2. DNA extraction and gene amplification

Genomic extractions were performed using the *ab*Genix™ automated DNA and RNA extraction system (AITbiotech Pte Ltd, Singapore), using the Animal Tissue Genomic DNA Extraction kits according to manufacturer’s instructions. All samples were processed separately.

We targeted the 313-bp region of cytochrome c oxidase subunit I (COI) gene because mini-barcodes were shown to perform just as well as the full-length barcodes for species-level identifications, and that loss in length for >200-bp barcodes did not result in demonstrable loss of information (Yeo et al., 2020). The MinION sequencing platform was benchmarked against Illumina NGS technology as the latter has already been validated to be just as accurate as Sanger sequencing (Baudhuin et al., 2015; Beck et al., 2016). We thus reasoned it was unnecessary to generate Sanger barcodes for comparison. PCR amplification was done using the mlCOIintF: 5’-GGWACWGGWTGAACWGTWTAYCCYCC-3’ (Leray et al., 2013) and LoboR1: 5’-TAAACYTCWGGRTGWCCRAARAAYCA-3’ (Lobo et al., 2013) primer combination. PCR primers were tagged with unique 8-bp barcode tags on the 5’ end to allow for downstream demultiplexing (Meier et al., 2016). Both the forward and reverse tags were unique to each specimen. Each PCR reaction comprised 2uL of template DNA, 2uL each of 10uM primer, 1uL of magnesium chloride, 1uL of bovine serum albumin (1mg/mL), 12.5uL of GoTaq Green Master Mix (Promega), and topped up to 25uL with sterile water. The thermal cycling profile used was as follows: 94°C for 60 secs; 5 cycles of 94°C for 30 secs, 48°C for 120 secs, 72°C for 60 secs, followed by 30 cycles of 94°C for 30 secs, 54°C for 120 secs, 72°C for 60 secs, and a final extension for 5 mins at 72°C. A subset of products were run on 2% agarose gels stained with GelRed (Cambridge Bioscience) to ensure amplification success, while all negative controls were screened to ensure they were clean.

We then pooled the PCR products into 9 separate pools; pooling was done by plate as we had up to 96 unique barcode tag combinations. The pools were then cleaned using 1.1× Sera-Mag™ Magnetic SpeedBeads™ (GE Healthcare Life Sciences) in 18% polyethylene glycol (PEG) buffer suspension:DNA ratio. The same purified amplicon pools were then used for both Illumina and MinION sequencing to allow for direct comparison of sequencing accuracy between platforms. Illumina sequencing technology, especially MiSeq, is known to be highly accurate (Loman et al., 2012; Fox et al., 2014), thereby serving as a suitable reference baseline to determine accuracy of MinION barcodes.

### 2.3. MinION barcoding

We prepared two different sets of amplicon pools for MinION sequencing. The first set comprised a single library pool with 96 amplicons, while the 8 remaining libraries made up the second set. About 300ng and 700ng of starting material were used for the first and second sets respectively. MinION libraries were prepared using the Ligation Sequencing Kit (SQK-LSK109), with the following modifications: (1) end-repair and dA-tailing reactions were incubated in a thermocycler at 20°C for 30 mins, followed by 65°C for 30 mins, and (2) ligation reactions were incubated in the thermocycler for 15 mins at 20°C. An additional multiplexing step was performed for the second set using the Native Barcoding Expansion (1-12; EXP-NDB-104), using the same modifications described above. Library preparation took approximately 2 h for the first set and 4–5 h for the second set. Sequencing was performed on two separate R9.4.1. flow cells on MinKNOW for Ubuntu 18 (v4.0.5), and each instance was run for 24 h.

### 2.4. Illumina barcoding

For Illumina-based validation, we prepared PCR-free libraries using NEBNext Ultra II DNA library prep kit (New England Biolabs), but with TruSeq DNA Single Indexes (Set B, Illumina), following the manufacturers’ instructions up till the adapter ligation step. Libraries were then cleaned using 1.1× Sera-Mag PEG suspension before final pooling in equimolar ratios and subsequent sequencing over one lane of Illumina MiSeq platform (251×251-bp) at the Genome Institute of Singapore.

### 2.5. Bioinformatics pipeline

For MinION reads, the raw fast5 files were uploaded onto a computer cluster for basecalling (guppy version 3.1.5+781ed57). For both datasets, no quality filtering criteria was applied during the basecalling process. For the second flow cell, *guppy_barcoder* was used to further demultiplex basecalled reads by native barcodes. The MinION reads were then analysed using the *miniBarcoder.py* script (Srivathsan et al., 2018). We performed the recommended full search (*-D 0*) on the dataset using the unique tag mode (*-m 1*), which permitted only 1-bp mismatch between tags (Srivathsan et al., 2018). Because our tags were shorter, we performed two different variations of the unique tag search; (i) ‘*full*’, and (ii) ‘*half*’. (see Supplementary Table 2). The latter setting was run to minimise the likelihood of erroneous demultiplexing by preventing binning of reads that had only mutant tags. Any resulting MAFFT barcode that had <10× read coverage and >1% of ambiguous bases called as Ns were also removed. For generating RACON barcodes, we adhered to the GraphMap v0.5.2 (Sović et al., 2016) max error rate of 0.15 that was suggested for 1D reads (Srivathsan et al., 2019). Both MAFFT and RACON barcodes were then subjected to amino acid correction (Srivathsan et al., 2018), using publicly available sequences on GenBank (*nt* database downloaded 16th July 2019) to yield MAFFT+AA and RACON+AA barcodes respectively. As our sample set consisted of fauna from different phyla, the appropriate genetic code (option-*g*) needed to be applied in the correction process. We used codes 2, 4, 5, and 9 for vertebrate, Cnidaria, invertebrate, and echinoderms and flatworm samples respectively. We also varied the namino option (Srivathsan et al., 2019). The final step was to align the amino acid corrected barcodes and perform consensus calling to derive consolidated barcodes.

For Illumina-based barcoding, we ran a modified bioinformatics pipeline from (Sze et al., 2018) and (Leveque et al., 2019). Briefly, paired-end reads were merged using PEAR v0.9.11 (Zhang et al., 2014) and OBITools v1.2.11 (Boyer et al., 2016) was used for demultiplexing and further downstream processing of assembled reads. All the steps were similar, except that instead of *obisplit*, we used *obisubset* to distribute the reads across samples. We applied the following quality filtering criteria to consider the derived Illumina barcodes as valid: (1) total reads assigned to a sample needed a minimum 10× coverage, and (2) if there were secondary reads assigned to that sample, the dominant read needed to be at least five times more abundant as the next most dominant sequence (Srivathsan et al., 2018). A translation check was then performed on Geneious R11 v11.1.5 (Kearse et al., 2012); this was to ensure our Illumina reference sequences were of mitochondrial origin.

### 2.6. MinION barcode accuracy and clustering congruence

As the primary aim of this paper was to showcase the feasibility of MinION-based barcoding in ARMS research, we focused our analysis on demonstrating the reliability of error-corrected MinION barcodes rather than drawing any ecological- or community-level inferences from our results.

Both the MinION and Illumina barcodes were screened for contamination using BLASTn (Camacho et al., 2009) against the same *nt* database used previously for error-correction of MinION barcodes. BLAST hits that had at least 70% BLAST match to the database and a minimum 250 bp overlap were parsed through readsidentifier v1.0 (Srivathsan et al., 2015). We used the MAFFT dataset as it was the largest barcode dataset, and retrospectively filtered the other datasets for contaminants. For our contamination check, we used the morpho-phylum classifications made during the sample vouchering process to match against taxonomic output from readsidentifier. Samples that failed this congruence check were followed-up with voucher examinations to check for potential wrong morpho-phylum assignments (i.e. pre-sorting error). If pre-sorting error was deemed unlikely, the barcode was subsequently removed from the dataset. Any barcode that matched a non-metazoan sequence (e.g. bacteria) was also excluded.

We assessed sequencing accuracy of our clean MinION barcodes with the Illumina reference barcodes using the *assess_corrbarcodes_wref.py* and *assess_uncorrbarcodes_wref.py* scripts; accuracy is defined as the number of perfect matches over the total number of bases compared (Srivathsan et al., 2018, 2019). Any MinION barcode that differed from its Illumina reference by >3% was deemed erroneous and removed; this was because the same amplicon pools were used, and hence, derived barcodes should be identical regardless of platform used. Any differences, if any, could indicate erroneous binning of MinION reads into the wrong sample (i.e. demultiplexing error).

We then determined whether the two sequencing approaches differed in the number of molecular operational taxonomic units (MOTUs) attained. Overlapping Illumina and MinION barcodes from each dataset were separately aligned on MAFFT v7.407 (Katoh and Standley, 2013) under default parameters. Objective clustering was carried out with gaps treated as missing data, grouping sequences into MOTUs based on uncorrected *p*-distances, testing thresholds of 2–4% (Meier et al., 2006; Srivathsan and Meier, 2012). We then performed a match ratio check to investigate if MinION clustered in the same way as Illumina barcodes (Ahrens et al., 2016; Srivathsan et al., 2019; Yeo et al., 2020).

### 2.7. Macrofaunal biodiversity of ARMS

We evaluated barcode accuracy and MOTU congruence performance across the MinION-generated datasets to select the best-performing consolidated barcode dataset for biodiversity analysis. Objective clustering was performed on the chosen dataset (in full) at 3% clustering threshold, with resultant MOTUs subjected to BLASTn. Only BLAST hits that had at least 80% match to the *nt* database, and had a minimum overlap of 250 bp parsed through readsidentifier to obtain taxonomic identities. We then performed a morphological examination of randomly-selected MOTUs to check if identities and MOTU members made sense.

### 2.8. Sequencing costs

We also calculated the sequencing costs associated with both MinION and Illumina to examine how they compared with the Sanger method. Our calculations did not consider the entry cost of sequencing hardware (i.e. the price of a MinION sequencer or Illumina MiSeq), which we assumed were readily accessible. For MinION sequencing, we assumed that reagents and flow cells were purchased separately because this provides a more meaningful gauge as the MinION starter pack is usually a one-time purchase for most users. For Illumina sequencing, we used publicly available sequencing costs (http://www.biotech.cornell.edu/brc/genomics/services/price-list#miseq).

## 3. Results

### 3.1. Macrofaunal fraction of ARMS

We obtained 725 specimens, representing six different phyla, from 12 ARMS units across four reef sites in Singapore. Samples were largely dominated by arthropods (314 samples), followed by molluscs (148 samples) and annelids (147 samples; Figure 2). In sum, 767 amplicons, comprising 725 specimens and 42 negative controls were sequenced on both Illumina and MinION platforms.

**Figure 2.**
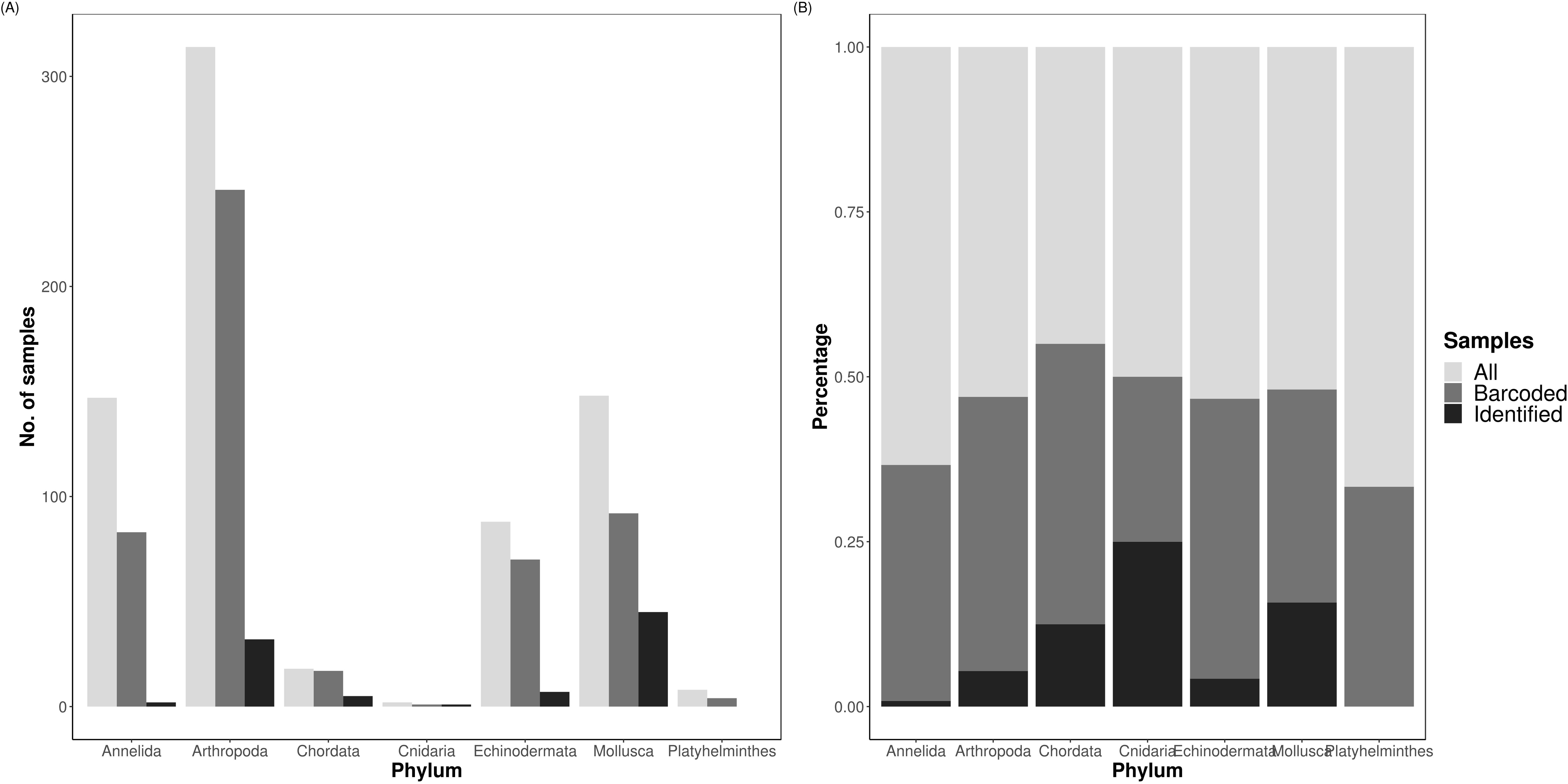
Total diversity obtained from ARMS across the different phyla by abundance (A) and proportion (B), showing the total number of samples obtained (“All”), number of barcodes obtained at the consolidated stage, namino=1 setting (“Barcoded”), and number of consolidated barcodes with species-level identification at ≥97% (“Identified”).

### 3.2. MinION barcoding

The first MinION flow cell generated 17,912,094 reads. As processing the entire dataset was computationally intensive, we subsampled the data at the 15-minute mark of sequencing to obtain the first 280,000 reads. The second flow cell had ceased after 3 h and generated 958,112 reads. Given that 98% of barcodes (or ~500 samples) could be obtained at 100× coverage within 2 h of sequencing (Srivathsan et al., 2018), we proceeded with downstream analysis as a 3-h run was expected to be sufficient for our sample size. The combined dataset comprised 1,238,112 raw reads, of which 1,091,301 reads remained after *guppy_barcoder* (Table 1).

**Table 1.**
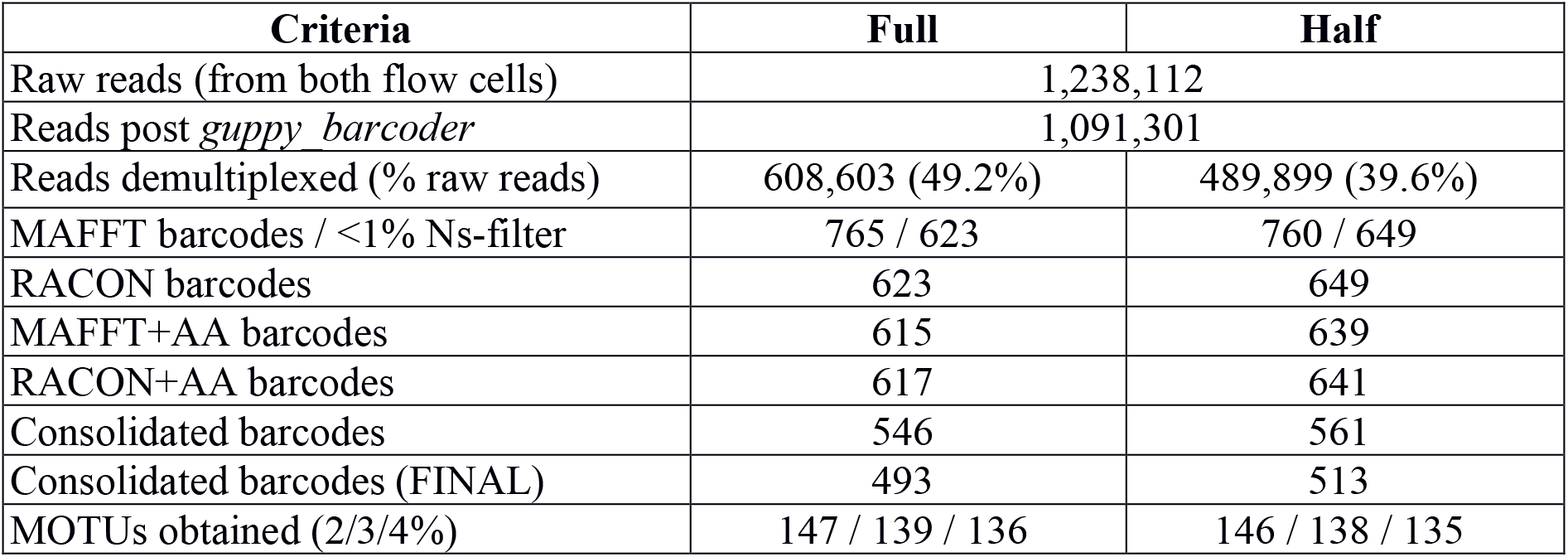
Reads and barcodes obtained across the different datasets. A total of 767 amplicons (of which 725 were ARMS samples) were barcoded. The amino acid corrected barcodes for MAFFT, RACON and eventual consolidated barcodes were obtained from namino=1 setting. Final refers to the final dataset with accepted barcodes, after removing contaminants.

| Criteria | Full | Half |
| --- | --- | --- |
| Raw reads (from both flow cells) | 1,238,112 |  |
| Reads post *guppy_barcode* | 1,091,301 |  |
| Reads demultiplexed (% raw reads) | 608,603 (49.2%) | 489,899 (39.6%) |
| MAFFT barcodes / <1% Ns-filter | 765 / 623 | 760 / 649 |
| RACON barcodes | 623 | 649 |
| MAFFT+AA barcodes | 615 | 639 |
| RACON+AA barcodes | 617 | 641 |
| Consolidated barcodes | 546 | 561 |
| Consolidated barcodes (FINAL) | 493 | 513 |
| MOTUs obtained (2/3/4%) | 147 / 139 / 136 | 146 / 138 / 135 |

We achieved 39 or 49% demultiplexing success depending on the criteria used (Table 1), with the ‘*full*’ and ‘*half*’ dataset capturing 608,603 and 489,899 reads respectively. Despite these differences, we obtained ~760 preliminary MAFFT barcodes across both datasets. Although the ‘*full*’ dataset yielded more preliminary MAFFT barcodes, it retained ~20 fewer barcodes compared to the ‘*half*’ dataset after filtering for >1% ambiguous bases. We then ran the entire *miniBarcoder* pipeline for all datasets, and found that while the ‘*half*’ dataset consistently had more barcodes than the ‘*full*’ dataset, MOTUs obtained were found to be highly similar (Table 1). We present results based on the ‘*half*’ dataset as it was the larger barcode dataset.

We obtained 649 MAFFT and RACON barcodes; 639 MAFFT+AA barcodes and 641 RACON+AA barcodes remained after amino-acid error correction; these corrected barcodes were then consolidated to obtain 561 consolidated barcodes (Table 1). All three types of error-corrected barcodes still retained a low number of ambiguities (coded as Ns), with MAFFT+AA barcodes having the most, and consolidated barcodes having the least ambiguities (Figure 3). Overall, MinION barcoding success was 77.4% (561 out of 725 specimens). None of the 42 negative controls passed the Ns-filter stage of the MAFFT barcode step. Of the 561 consolidated barcodes, a further 48 barcodes were removed because they failed the morpho-phylum and barcode congruence check. Failure was attributed to cross-contamination after confirmatory checks with the vouchers. Notably, the same 48 samples failed on the Illumina platform. No barcode was found to have top BLAST hit to non-metazoan sequences. We retained 513–514 consolidated barcodes for further analysis.

**Figure 3.**
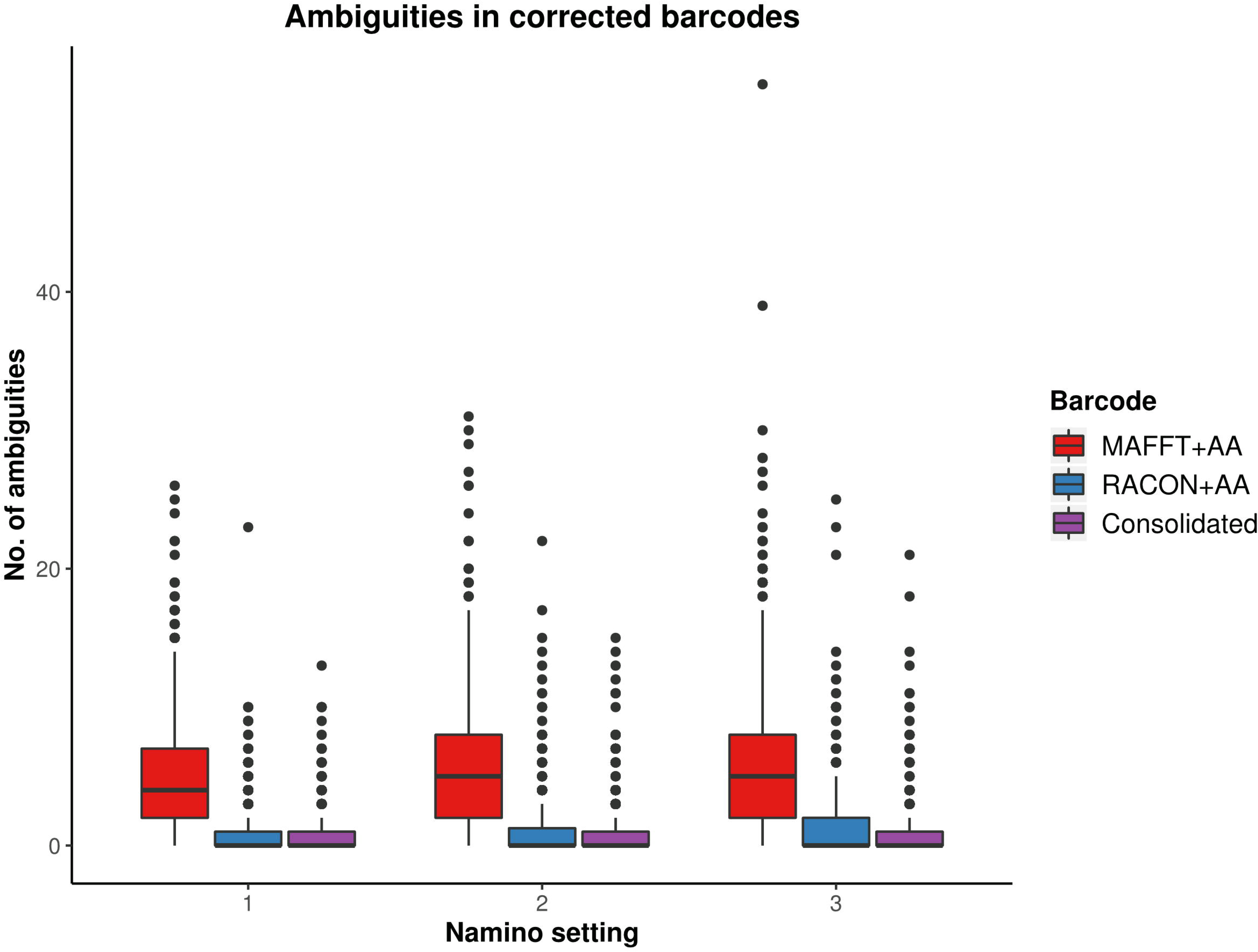
Ambiguities remaining in the three types of error-corrected MinION barcodes: MAFFTT+AA (red), RACON+AA (blue), and consolidated (purple) across varying namino settings (1−3).

### 3.3. Comparing MinION and Illumina barcodes

For our Illumina reference barcodes, 21,854,748 reads were generated on one MiSeq lane; 645 barcodes remained after applying sequencing filters. Five barcodes failed the translation check and were removed; 640 Illumina reference barcodes remained.

In terms of sequencing accuracy, we found all MinION datasets had near-perfect barcode accuracy when compared with Illumina barcodes, scoring at least 99.95% for accuracy; with the MAFFT dataset achieving 100% accuracy (Table 2). In addition, no MinION barcode had >3% mismatch when evaluated against the Illumina reference. MAFFT barcodes generally had few to no mismatches, while RACON barcodes were marginally less accurate than MAFFT barcodes (Table 2). The uncorrected barcode datasets scored higher on accuracy, but a fair proportion of them (>100 samples for each barcode dataset) had internal gaps. The error-correction pipeline was able to resolve this issue such that only 6–13 samples had gaps. It was the consolidated datasets that yielded the most barcodes with no gaps and the fewest mismatches (Table 2).

**Table 2.** Accuracy and gaps observed when MinION™ barcodes from the ‘*half*’ dataset were compared with Illumina barcodes. Accuracy is defined as the number of perfect matches across total number of base pairs compared, expressed in percentage (%).

| Barcode | No. of overlapping barcodes compared | Accuracy of MinION™ barcodes (%) | Number of samples with internal gaps |
| --- | --- | --- | --- |
| MAFFT Ns-filter | 584 | 100.000 | 520 |
| RACON | 584 | 99.994 | 186 |
| MAFFT+AA (namino=1) | 576 | 99.958 | 13 |
| MAFFT+AA (namino=2) | 576 | 99.971 | 12 |
| MAFFT+AA (namino=3) | 576 | 99.971 | 13 |
| RACON+AA (namino=1) | 577 | 99.967 | 6 |
| RACON+AA (namino=2) | 577 | 99.975 | 7 |
| RACON+AA (namino=3) | 577 | 99.972 | 6 |
| Consolidated (namino=1) | 507 | 99.989 | 3 |
| Consolidated (namino=2) | 507 | 99.989 | 3 |
| Consolidated (namino=3) | 508 | 99.987 | 3 |

The MOTUs obtained from objective clustering for MinION and Illumina barcode datasets were highly congruent, differing by up to just 4 MOTUs depending on the dataset (Table 3). MOTUs obtained were also fairly stable across the 2–4% *p*-distance thresholds tested for both sequencing datasets; match ratios were also very high (≥0.96) across the datasets (Table 3). Again, the consolidated barcodes performed the best, with the same number of MOTUs obtained, and perfect match ratios.

**Table 3.** The number of molecular operational taxonomic units (MOTUs) obtained for each overlapping dataset at 2–4% threshold, and the differences in MOTUs between MinION (’*half*’ dataset) and Illumina MOTUs.

| Barcode | No. of barcodes | MOTUs obtained (MinION/ Illumina) |  |  | MOTU Difference (MinION–Illumina) |  |  | Match Ratio |  |  |
| --- | --- | --- | --- | --- | --- | --- | --- | --- | --- | --- |
|  |  | 2% | 3% | 4% | 2% | 3% | 4% | 2% | 3% | 4% |
| MAFFT Ns-filter | 584 | 163/164 | 157/156 | 152/151 | -1 | 1 | 1 | 0.979 | 0.971 | 0.990 |
| RACON | 584 | 162/160 | 153/156 | 150/151 | 2 | -3 | -1 | 0.981 | 0.977 | 0.990 |
| MAFFT+AA (namino=1) | 576 | 158/160 | 151/152 | 147/148 | -2 | -1 | -1 | 0.981 | 0.990 | 0.990 |
| MAFFT+AA (namino=2) | 576 | 158/160 | 151/152 | 147/148 | -2 | -1 | -1 | 0.981 | 0.990 | 0.990 |
| MAFFT+AA (namino=3) | 576 | 158/160 | 151/152 | 147/148 | -2 | -1 | -1 | 0.981 | 0.990 | 0.990 |
| RACON+AA (namino=1) | 577 | 156/160 | 152/152 | 148/148 | -4 | 0 | 0 | 0.962 | 1.000 | 1.000 |
| RACON+AA (namino=2) | 577 | 161/160 | 152/152 | 148/148 | 1 | 0 | 0 | 0.990 | 1.000 | 1.000 |
| RACON+AA (namino=3) | 577 | 160/160 | 152/152 | 148/148 | 0 | 0 | 0 | 0.981 | 1.000 | 1.000 |
| Consolidated (namino=1) | 507 | 144/144 | 136/136 | 133/133 | 0 | 0 | 0 | 1.000 | 1.000 | 1.000 |
| Consolidated (namino=2) | 507 | 144/144 | 136/136 | 133/133 | 0 | 0 | 0 | 1.000 | 1.000 | 1.000 |
| Consolidated (namino=3) | 508 | 143/144 | 136/136 | 133/133 | -1 | 0 | 0 | 0.990 | 1.000 | 1.000 |

We ultimately selected the consolidated barcodes (namino=1) dataset for biological analysis, taking into consideration that it was highly accurate (99.989%), had the least samples with internal gaps, no differences in MOTUs obtained when compared to Illumina references, and had perfect match ratios across different clustering thresholds (Tables 2 and 3). While the namino=2 consolidated barcode dataset performed similarly well, the namino=1 dataset was chosen because it contained less ambiguities (Figure 3). We obtained 513 clean MinION barcodes with the namino=1 consolidated barcode dataset, representing 138 MOTUs at 3% objective clustering threshold (Table 1). Only 26 of the 138 MOTUs could be delimited to species at ≥97% sequence similarity, of which 17 had species-level identification. Molluscan samples (6 MOTUs) had the highest identification success to species-level, followed by Arthropoda (5 MOTUs; Figure 2). A similar pattern of identification success was noted previously by Ip et al. (2019).

### 3.4. Sequencing costs

Overall, the per sample costs for both Illumina and MinION sequencing was found to be six-fold cheaper than the Sanger method (Table 4), costing an estimated US$3 per barcode. MinION barcodes were marginally more expensive than Illumina barcodes (US$3.27 vs. US$3.15) in this study. The higher cost was attributable to the additional flow cell and reagents used in this study; had we not used them, the MinION barcodes would have cost US$1.91 (Table 4).

**Table 4.** Cost comparison between MinION, Illumina and Sanger sequencing for this study. Publicly available pricing was used for Illumina sequencing costs (http://www.biotech.cornell.edu/brc/genomics/services/price-list#miseq). Cost for DNA extraction or PCR amplification is not reflected in this table because the steps and costs are the same.

| Sequencing | Package Cost (US\$) | Per Sample Cost (US\$) | Total (US\$) |
| --- | --- | --- | --- |
| MinION | \$900 per flow cell (used 2)<br>\$599 per Ligation Sequencing Kit (6 preps, used 2)<br>\$288 per Native Barcoding Expansion (6 preps, used 1)<br>\$1,170.24 per NEBNext ONT Companion Kit (24 preps, used 9)<br>\$1,275.53 for 60ml Agencourt AMPure XP | \$3.24 (767 amplicons in total)<br>*\$1.88 (if only one flow cell, one Ligation Sequencing reaction used, without Native Barcoding Expansion)*<br>\$0.03 (estimated from using ~130µl per library of 96 samples) | \$3.27 (\$1.91) |
| Illumina | \$2,121 for MiSeq sequencing cost<br>\$146.28 for TruSeq Set B adapters (48 libraries, used 9)<br>\$750.48 per NEBNext Ultra II Library Prep Kit for Illumina (24 preps, used 9)<br>\$1,275.53 for 60ml Agencourt AMPure XP | \$3.12 (767 amplicons in total)<br>\$0.03 (estimated from using ~130µl per library of 96 samples) | \$3.15 |
| Sanger | \$12,194 per Big Dye Terminator v3.1 Cycling Kit (1,000 reactions)<br>\$337.90 for 96-well plates (100 units)<br>\$4.68 per primer (100µM; approximately 500 reactions)<br>\$1187.60 for 50ml Aline Biosciences PureSEQ (10,000 reactions)<br>\$1.22 per well sequencing cost | \$24.40 (2 reactions for 1 sample)<br>\$0.07 (2 reactions for 1 sample)<br>\$0.02 (2 reactions for 1 sample)<br>\$0.24 (2 reactions for 1 sample)<br>\$2.45 (2 reactions for 1 sample) | \$27.18 |

## 4. Discussion

### 4.1. Cost- and time-savings with MinION-based barcoding

We here demonstrate a successful application of the MinION-based pipeline to process >2-mm samples from ARMS units deployed on tropical coral reefs. Our strategy is advantageous to researchers using ARMS due to cost and time savings, as well as the low outlay of sequencing equipment.

Most importantly, the costs involved in MinION sequencing are inversely related to the sample size. The abundance of organisms in the >2-mm fraction ranges from several hundreds to a thousand (Supplementary Table 1). Previously, such sample sizes would have been too small to be multiplexed cost-effectively on an Illumina or PacBio flow cell—one would need >10,000 specimens with Illumina NGS barcoding for US$1 barcodes (Meier et al., 2016; Srivathsan et al., 2018)—but large enough that Sanger sequencing becomes laborious and cost-prohibitive. MinION-barcoding is ideal for ARMS researchers because it caters to projects with small to moderate sample sizes. For example, Srivathsan et al. (2019) demonstrated sequencing costs as low as <US$0.35 per sample for ~3,500 specimens on a single flow cell. In this study, our sample size (725) using the MinION barcode costs more at US$3 per sample, but would be considerably cheaper (US$1.91 per sample) had we used a single flow cell (Table 4). Nevertheless, MinION barcoding (and even Illumina barcoding) remains considerably cheaper than Sanger barcoding (≥US$18 per barcode; Table 4; Meier et al., 2016). Furthermore, the ability to multiplex numerous samples on a flow cell saves considerable time. In this study, we obtained ~700 preliminary barcodes in 3.5 h, whereas the Sanger method for the equivalent sample size would take ~18 h. This is attributable to the laborious preparatory bench work of the latter, where sample pooling is not possible, and samples have to be processed in batches. By employing a multiplexing strategy in MinION barcoding, all the amplicons can be processed into a single library at one time. Overall, MinION barcoding is cheaper and faster for processing a larger number of samples than the present workflow for the ARMS >2-mm fraction.

Relatedly, the MinION hardware is also comparatively cheaper to obtain than most sequencers (e.g. ABI capillary or Illumina), with a starter pack costing approximately US$1,000. Most molecular laboratories would already have access to basic laboratory equipment (e.g. thermocyclers), so there is no additional hardware required to perform MinION barcoding. Moreover, the minimal computational prerequisites for the *miniBarcoder* pipeline further enhance the attractiveness of MinION barcoding. We completed the entire analysis pipeline on a 4-core computer (32 Gb RAM) within one week, meaning that a conventional laptop or desktop is sufficient. Additionally, the plug-and-play nature of the software makes it intuitive for users, who need only basic proficiency in the Ubuntu environment.

### 4.2. MinION barcodes are viable

Furthermore, the *miniBarcoder* pipeline affords flexibility while remaining scientifically robust. First, the *miniBarcoder* pipeline was able to accommodate our use of shorter tags (8 bp vs. 13 bp); the fact that none of our MinION barcodes differed by >3% from their Illumina references suggests high demultiplexing success, and that any failures in morpho-phylum and barcode congruence checks on both platforms were due to wet-lab errors. Notably, our second flow cell had two levels of multiplexing; the first via tagged primers, and the second via ONT native barcodes, which may increase margins for demultiplexing error, but we did not observe this occurring with our dataset, further demonstrating high demultiplexing fidelity. Second, we successfully used different settings during the unique tag search to increase barcode recovery (Supplementary Table 2). Despite lower demultiplexing success, we recovered a higher number of MAFFT N-filtered barcodes from the ‘*half*’ rather than ‘*full*’ dataset (Table 1). We posit that accepting mutant tags introduced more erroneous reads per sample in the ‘*full*’ dataset, which increased the number of ambiguous bases during consensus calling step when generating preliminary MAFFT barcodes and led to their subsequent removal. This suggests that read quality rather than numbers of demultiplexed reads is more important for recovering barcodes. We thus recommend that users toggle between settings to maximise barcode retrieval. Such tests do not take long and would translate to cost savings by reducing the need for re-sequencing. Third, we successfully applied the barcoding pipeline to a broad spectrum of >2-mm specimens, representing six different phyla (Figure 2), and managed to recover 513 consolidated MinION barcodes. This was possible through the careful application of the appropriate mitochondrial genetic code in the error-correction step. Our amplification and identification success rates were similar to a previous barcoding study by Ip et al. (2019), which employed a combination of Sanger and Illumina barcoding, further suggesting that MinION barcodes are viable and that performance success is defined more by primer choice rather than sequencing platform.

Ultimately, we opted for a MinION barcoding approach for the >2-mm size fraction over bulk-sample metabarcoding (as with the other size fractions) in order to remain true to the intended design of ARMS (Leray and Knowlton, 2015), which is to retain sequence-to-sample association. Such an approach will be helpful for the confirmation of previous species records, as demonstrated in other studies (Huang et al., 2014; Poquita-Du et al., 2017, 2019; Yip et al., 2018; Ng et al., 2019; Oh et al., 2019) and voucher-sequence matching would facilitate downstream integrative taxonomic work to either resolve cryptic species complexes (Bickford et al., 2007; Chang et al., 2018; Chan et al., 2019), or even describe new species (Srivathsan et al., 2019). The fact that our samples remain largely unidentifiable (only 12% of MOTUs could be identified to species at ≥97% sequence similarity) further impresses the need for postliminary taxonomic work to establish species-level identity—whether they are existing or new species—and add new information to DNA barcode repositories (e.g. Kutty et al., 2018; Ip et al., 2019). We emphasize that many of our samples remain unidentified due to lack of robust online database matches rather than the ambiguities that persist in the MinION barcodes after error-correction (Figure 3); the latter of which has been demonstrated to be of minor concern (Srivathsan et al., 2018, 2019; Ho et al., 2020). As nanopore sequencing chemistry continues to improve with the release of newer flow cells, it would be interesting to investigate how the recently-released R10.3 flow cell, which purportedly produces more accurate reads, fares in MinION barcoding.

It presently remains untested whether MinION sequencing can be applied to metabarcoding of the other size fractions. The *miniBarcoder* has been successfully applied to mixed food items, essentially small-scaled metabarcoding samples, but the reads had to be first parsed by identity and then manually grouped before consensus calling to obtain MAFFT barcodes (Ho et al., 2020). This is achievable in food items where taxa are limited and known, but likely unattainable for ARMS samples with the current *miniBarcoder* pipeline since the entire tree of life is represented, making manual grouping of reads arduous. It also becomes more challenging to group reads when it is nearly impossible to ascertain if reads are different due to sequencing error, or if they are truly biologically divergent. As such, we do not recommend the MinION for metabarcoding using R9.4.1 flow cells, though with aforementioned improvements in flow cell chemistry, and the release of high-throughput sequencers like GridION and PromethION, nanopore metabarcoding for ARMS research may be accomplished in the near future.

## 5. Conclusion

We here proposed MinION-based barcoding as an alternative sequencing approach to the initial pipeline (Leray and Knowlton, 2015) for rapid processing of the >2-mm size fraction involving hundreds to thousands of specimens. We have demonstrated that MinION-based barcoding is viable and highly accurate, by comparing our MinION barcodes to Illumina references generated from the same library pools. The few ambiguities that persist in the error-corrected MinION barcodes do not compromise biological inferences. MinION barcodes also exhibit congruent clustering patterns with reference barcodes, are cheaper than the existing Sanger barcodes, and an entire dataset of ~700 barcodes can be easily obtained in under 4 h. We conclude that this method would most certainly streamline the workflow for processing ARMS samples. The faster we are able to generate barcodes, the sooner the integrative taxonomic work and ecological analyses of marine biodiversity can begin.

## Authors’ Contributions

All authors collectively conceived the idea for the study and led the fieldwork. AI and MC vouchered the specimens, assisted by DH. AI performed the DNA extractions, gene amplifications, Illumina library preparation and data analysis. MC performed the MinION sequencing and data analysis. MC wrote the manuscript, with help from the other authors. All authors approved the final version of the manuscript for publication.

## Funding

Funding for this research was provided by the Singapore Ministry of Education Academic Research Fund Tier 1 (R-154-000-A63-114), the National Research Foundation, Prime Minister’s Office, Singapore under its Marine Science R&D Programme (MSRDP-P03), and an AXA postdoctoral fellowship to AB (R-154-000-649-507).

## Conflict of Interest

The authors declare that the research was conducted in the absence of any commercial or financial relationships that could be construed as a potential conflict of interest.

## Acknowledgements

We are grateful to the following for their help with fieldwork and processing: Yong Kit Samuel Chan, Zheng Bin Randolph Quek, Sudhanshi Sanjeev Jain, Ren Min Oh, Jovena Chun Ling Seah, Zack Chen, Chin Soon Lionel Ng, Joy Shu Yee Wong, Sherlyn Sher Qing Lim, Zhi Ting Yip and Jun Wei Phua. We also thank Amrita Srivathsan for her advice on the *miniBarcoder* pipeline, and National Supercomputing Centre, Singapore (https://www.nscc.sg), for use of their computational resources.

## References

Ahrens, D., Fujisawa, T., Krammer, H.-J., Eberle, J., Fabrizi, S., and Vogler, A. P. (2016). Rarity and Incomplete Sampling in DNA-Based Species Delimitation. Syst. Biol. 65, 478–494. doi:10.1093/sysbio/syw002.

Al-Rshaidat, M. M. D., Snider, A., Rosebraugh, S., Devine, A. M., Devine, T. D., Plaisance, L., et al. (2016). Deep COI sequencing of standardized benthic samples unveils overlooked diversity of Jordanian coral reefs in the northern Red Sea. Genome 59, 724–737. doi:10.1139/gen-2015-0208.

Appeltans, W., Ahyong, S. T., Anderson, G., Angel, M. V., Artois, T., Bailly, N., et al. (2012). The magnitude of global marine species diversity. Current Biology 22, 2189–2202. doi:10.1016/j.cub.2012.09.036.

Baudhuin, L. M., Lagerstedt, S. A., Klee, E. W., Fadra, N., Oglesbee, D., and Ferber, M. J. (2015). Confirming Variants in Next-Generation Sequencing Panel Testing by Sanger Sequencing. J. Mol. Diagn. 17, 456–461. doi:10.1016/j.jmoldx.2015.03.004.

Beck, T. F., Mullikin, J. C., NISC Comparative Sequencing Program, and Biesecker, L. G. (2016). Systematic Evaluation of Sanger Validation of Next-Generation Sequencing Variants. Clin. Chem. 62, 647–654. doi:10.1373/clinchem.2015.249623.

Bickford, D., Lohman, D. J., Sodhi, N. S., Ng, P. K. L., Meier, R., Winker, K., et al. (2007). Cryptic species as a window on diversity and conservation. Trends Ecol. Evol. 22, 148–155. doi:10.1016/j.tree.2006.11.004.

Boyer, F., Mercier, C., Bonin, A., Le Bras, Y., Taberlet, P., and Coissac, E. (2016). obitools: a unix-inspired software package for DNA metabarcoding. Mol. Ecol. Resour. 16, 176–182. doi:10.1111/1755-0998.12428.

Camacho, C., Coulouris, G., Avagyan, V., Ma, N., Papadopoulos, J., Bealer, K., et al. (2009). BLAST : architecture and applications. BMC Bioinformatics 10, 421. doi:10.1186/1471-2105-10-421.

Carvalho, S., Aylagas, E., Villalobos, R., Kattan, Y., Berumen, M., and Pearman, J. K. (2019). Beyond the visual: using metabarcoding to characterize the hidden reef cryptobiome. Proc. Biol. Sci. 286, 20182697. doi:10.1098/rspb.2018.2697.

Chang, J. J. M., Tay, Y. C., Ang, H. P., Tun, K. P. P., Chou, L. M., Meier, R., et al. (2018). Molecular and anatomical analyses reveal that Peronia verruculata (Gastropoda: Onchidiidae) is a cryptic species complex. Contributions to Zoology 87, 149–165. doi:10.1163/18759866-08703002.

Chan, I. Z. W., Chang, J. J. M., Huang, D., and Todd, P. A. (2019). Colour pattern measurements successfully differentiate two cryptic Onchidiidae Rafinesque, 1815 species. Marine Biodiversity 49, 1743–1750. doi:10.1007/s12526-019-00940-4.

Costello, M. J., and Wilson, S. P. (2011). Predicting the number of known and unknown species in European seas using rates of description. Global Ecology and Biogeography 20, 319–330. doi:10.1111/j.1466-8238.2010.00603.x.

Danovaro, R., Carugati, L., Berzano, M., Cahill, A. E., Carvalho, S., Chenuil, A., et al. (2016). Implementing and Innovating Marine Monitoring Approaches for Assessing Marine Environmental Status. Frontiers in Marine Science 3. doi:10.3389/fmars.2016.00213.

David, R., Uyarra, M. C., Carvalho, S., Anlauf, H., Borja, A., Cahill, A. E., et al. (2019). Lessons from photo analyses of Autonomous Reef Monitoring Structures as tools to detect (bio-)geographical, spatial, and environmental effects. Mar. Pollut. Bull. 141, 420–429. doi:10.1016/j.marpolbul.2019.02.066.

Fox, E. J., Reid-Bayliss, K. S., Emond, M. J., and Loeb, L. A. (2014). Accuracy of Next Generation Sequencing Platforms. Next Gener Seq Appl 1. doi:10.4172/jngsa.1000106.

Glenn, T. C. (2011). Field guide to next-generation DNA sequencers. Mol. Ecol. Resour. 11, 759–769. doi:10.1111/j.1755-0998.2011.03024.x.

Hayes, K. R., Cannon, R., Neil, K., and Inglis, G. (2005). Sensitivity and cost considerations for the detection and eradication of marine pests in ports. Mar. Pollut. Bull. 50, 823–834. doi:10.1016/j.marpolbul.2005.02.032.

Hazeri, G., Rahayu, D. L., Subhan, B., Sembiring, A., Anggoro, A. W., Ghozali, A. T., et al. (2019). Latitudinal species diversity and density of cryptic crustacean (Brachyura and Anomura) in micro-habitat Autonomous Reef Monitoring Structures across Kepulauan Seribu, Indonesia. Biodiversitas 20. doi:10.13057/biodiv/d200540.

Ho, J. K. I., Puniamoorthy, J., Srivathsan, A., and Meier, R. (2020). MinION sequencing of seafood in Singapore reveals creatively labelled flatfishes, confused roe, pig DNA in squid balls, and phantom crustaceans. Food Control. doi:10.1101/826032.

Huang, D., Cranston, P. S., and Cheng, L. (2014). A complete species phylogeny of the marine midge Pontomyia (Diptera:Chironomidae) reveals a cosmopolitan species and a new synonym. Invertebrate Systematics 28, 277. doi:10.1071/is13059.

Hurley, K. K. C., Timmers, M. A., Scott Godwin, L., Copus, J. M., Skillings, D. J., and Toonen, R. J. (2016). An assessment of shallow and mesophotic reef brachyuran crab assemblages on the south shore of O‘ahu, Hawai‘i. Coral Reefs 35, 103–112. doi:10.1007/s00338-015-1382-z.

Ip, Y. C. A., Tay, Y. C., Gan, S. X., Ang, H. P., Tun, K., Chou, L. M., et al. (2019). From marine park to future genomic observatory? Enhancing marine biodiversity assessments using a biocode approach. Biodiversity Data Journal 7, e46833. doi:10.3897/BDJ.7.e46833.

Katoh, K., and Standley, D. M. (2013). MAFFT multiple sequence alignment software version 7: improvements in performance and usability. Mol. Biol. Evol. 30, 772–780. doi:10.1093/molbev/mst010.

Kearse, M., Moir, R., Wilson, A., Stones-Havas, S., Cheung, M., Sturrock, S., et al. (2012). Geneious Basic: an integrated and extendable desktop software platform for the organization and analysis of sequence data. Bioinformatics 28, 1647–1649. doi:10.1093/bioinformatics/bts199.

Knowlton, N., Brainard, R. E., Fisher, R., Moews, M., Plaisance, L., and Julian Caley, M. (2010). “Coral Reef Biodiversity,” in Life in the World’s Oceans: Diversity, Distribution, and Abundance, ed. A. D. McIntyre, 65–78. doi:10.1002/9781444325508.ch4.

Krehenwinkel, H., Pomerantz, A., Henderson, J. B., Kennedy, S. R., Lim, J. Y., Swamy, V., et al. (2019). Nanopore sequencing of long ribosomal DNA amplicons enables portable and simple biodiversity assessments with high phylogenetic resolution across broad taxonomic scale. Gigascience 8. doi:10.1093/gigascience/giz006.

Kutty, S. N., Wang, W., Ang, Y. C., Tay, Y. C., Ho, J. K. I., and Meier, R. (2018). Next-Generation identification tools for Nee Soon freshwater swamp forest, Singapore. Gardens’ Bulletin Singapore 70, 155–173. doi:10.26492/gbs70(suppl.1).2018-08.

Leray, M., and Knowlton, N. (2015). DNA barcoding and metabarcoding of standardized samples reveal patterns of marine benthic diversity. Proc. Natl. Acad. Sci. U. S. A. 112, 2076–2081. doi:10.1073/pnas.1424997112.

Leray, M., Yang, J. Y., Meyer, C. P., Mills, S. C., Agudelo, N., Ranwez, V., et al. (2013). A new versatile primer set targeting a short fragment of the mitochondrial COI region for metabarcoding metazoan diversity: application for characterizing coral reef fish gut contents. Front. Zool. 10, 34. doi:10.1186/1742-9994-10-34.

Leveque, S., Afiq-Rosli, L., Ip, Y. C. A., Jain, S. S., and Huang, D. (2019). Searching for phylogenetic patterns of Symbiodiniaceae community structure among Indo-Pacific Merulinidae corals. PeerJ 7, e7669. doi:10.7717/peerj.7669.

Lobo, J., Costa, P. M., Teixeira, M. A. L., Ferreira, M. S. G., Costa, M. H., and Costa, F. O. (2013). Enhanced primers for amplification of DNA barcodes from a broad range of marine metazoans. BMC Ecol. 13, 34. doi:10.1186/1472-6785-13-34.

Loman, N. J., Misra, R. V., Dallman, T. J., Constantinidou, C., Gharbia, S. E., Wain, J., et al. (2012). Performance comparison of benchtop high-throughput sequencing platforms. Nat. Biotechnol. 30, 434–439. doi:10.1038/nbt.2198.

Lu, H., Giordano, F., and Ning, Z. (2016). Oxford Nanopore MinION Sequencing and Genome Assembly. Genomics, Proteomics & Bioinformatics 14, 265–279. doi:10.1016/j.gpb.2016.05.004.

Maestri, S., Cosentino, E., Paterno, M., Freitag, H., Garces, J. M., Marcolungo, L., et al. (2019). A Rapid and Accurate MinION-Based Workflow for Tracking Species Biodiversity in the Field. Genes 10. doi:10.3390/genes10060468.

Meier, R., Shiyang, K., Vaidya, G., and Ng, P. K. L. (2006). DNA Barcoding and Taxonomy in Diptera: A Tale of High Intraspecific Variability and Low Identification Success. Systematic Biology 55, 715–728. doi:10.1080/10635150600969864.

Meier, R., Wong, W., Srivathsan, A., and Foo, M. (2016). $1 DNA barcodes for reconstructing complex phenomes and finding rare species in specimen-rich samples. Cladistics 32, 100–110. doi:10.1111/cla.12115.

Mikheyev, A. S., and Tin, M. M. Y. (2014). A first look at the Oxford Nanopore MinION sequencer. Molecular Ecology Resources 14, 1097–1102. doi:10.1111/1755-0998.12324.

Mora, C., Tittensor, D. P., Adl, S., Simpson, A. G. B., and Worm, B. (2011). How many species are there on Earth and in the ocean? PLoS Biol. 9, e1001127. doi:10.1371/journal.pbio.1001127.

Ng, C. S. L., Jain, S. S., Nguyen, N. T. H., Sam, S. Q., Kikuzawa, Y. P., Chou, L. M., et al. (2019). New genus and species record of reef coral Micromussa amakusensis in the southern South China Sea. Marine Biodiversity Records 12. doi:10.1186/s41200-019-0176-3.

Oh, R. M., Neo, M. L., Yap, N. W. L., Jain, S. S., Tan, R., Chen, C. A., et al. (2019). Citizen science meets integrated taxonomy to uncover the diversity and distribution of Corallimorpharia in Singapore. Raffles Bulletin of Zoology 67, 306–321. doi:10.26107/RBZ-2019-0022.

Pearman, J. K., Anlauf, H., Irigoien, X., and Carvalho, S. (2016). Please mind the gap – Visual census and cryptic biodiversity assessment at central Red Sea coral reefs. Marine Environmental Research 118, 20–30. doi:10.1016/j.marenvres.2016.04.011.

Pearman, J. K., Aylagas, E., Voolstra, C. R., Anlauf, H., Villalobos, R., and Carvalho, S. (2019). Disentangling the complex microbial community of coral reefs using standardized Autonomous Reef Monitoring Structures (ARMS). Mol. Ecol. 28, 3496–3507. doi:10.1111/mec.15167.

Pearman, J. K., Leray, M., Villalobos, R., Machida, R. J., Berumen, M. L., Knowlton, N., et al. (2018). Cross-shelf investigation of coral reef cryptic benthic organisms reveals diversity patterns of the hidden majority. Sci. Rep. 8, 8090. doi:10.1038/s41598-018-26332-5.

Pennesi, C., and Danovaro, R. (2017). Assessing marine environmental status through microphytobenthos assemblages colonizing the Autonomous Reef Monitoring Structures (ARMS) and their potential in coastal marine restoration. Mar. Pollut. Bull. 125, 56–65. doi:10.1016/j.marpolbul.2017.08.001.

Plaisance, L., Brainard, R., Julian Caley, M., and Knowlton, N. (2011a). Using DNA Barcoding and Standardized Sampling to Compare Geographic and Habitat Differentiation of Crustaceans: A Hawaiian Islands Example. Diversity 3, 581–591. doi:10.3390/d3040581.

Plaisance, L., Caley, M. J., Brainard, R. E., and Knowlton, N. (2011b). The diversity of coral reefs: what are we missing? PLoS One 6, e25026. doi:10.1371/journal.pone.0025026.

Poquita-Du, R. C., Quek, Z. B. R., Jain, S. S., Schmidt-Roach, S., Tun, K., Heery, E. C., et al. (2019). Last species standing: loss of Pocilloporidae corals associated with coastal urbanization in a tropical city state. Marine Biodiversity 49, 1727–1741. doi:10.1007/s12526-019-00939-x.

Poquita-Du, R., Ng, C. S. L., Loo, J. B., Afiq-Rosli, L., Tay, Y. C., Todd, P., et al. (2017). New evidence shows that Pocillopora “damicornis-like” corals in Singapore are actually Pocillopora acuta (Scleractinia: Pocilloporidae). Biodiversity Data Journal 5, e11407. doi:10.3897/bdj.5.e11407.

Quail, M. A., Smith, M., Coupland, P., Otto, T. D., Harris, S. R., Connor, T. R., et al. (2012). A tale of three next generation sequencing platforms: comparison of Ion Torrent, Pacific Biosciences and Illumina MiSeq sequencers. BMC Genomics 13, 341. doi:10.1186/1471-2164-13-341.

Ransome, E., Geller, J. B., Timmers, M., Leray, M., Mahardini, A., Sembiring, A., et al. (2017). The importance of standardization for biodiversity comparisons: A case study using autonomous reef monitoring structures (ARMS) and metabarcoding to measure cryptic diversity on Mo’orea coral reefs, French Polynesia. PLOS ONE 12, e0175066. doi:10.1371/journal.pone.0175066.

Shokralla, S., Gibson, J. F., Nikbakht, H., Janzen, D. H., Hallwachs, W., and Hajibabaei, M. (2014). Next-generation DNA barcoding: using next-generation sequencing to enhance and accelerate DNA barcode capture from single specimens. Mol. Ecol. Resour. 14, 892–901. doi:10.1111/1755-0998.12236.

Sović, I., Šikić, M., Wilm, A., Fenlon, S. N., Chen, S., and Nagarajan, N. (2016). Fast and sensitive mapping of nanopore sequencing reads with GraphMap. Nat. Commun. 7, 11307. doi:10.1038/ncomms11307.

Srivathsan, A., Baloğlu, B., Wang, W., Tan, W. X., Bertrand, D., Ng, A. H. Q., et al. (2018). A MinION-based pipeline for fast and cost-effective DNA barcoding. Molecular Ecology Resources 18, 1035–1049. doi:10.1111/1755-0998.12890.

Srivathsan, A., Hartop, E., Puniamoorthy, J., Lee, W. T., Kutty, S. N., Kurina, O., et al. (2019). Rapid, large-scale species discovery in hyperdiverse taxa using 1D MinION sequencing. BMC Biology 17. doi:10.1101/622365.

Srivathsan, A., and Meier, R. (2012). On the inappropriate use of Kimura-2-parameter (K2P) divergences in the DNA-barcoding literature. Cladistics 28, 190–194. doi:10.1111/j.1096-0031.2011.00370.x.

Srivathsan, A., Sha, J. C. M., Vogler, A. P., and Meier, R. (2015). Comparing the effectiveness of metagenomics and metabarcoding for diet analysis of a leaf-feeding monkey (Pygathrix nemaeus). Mol. Ecol. Resour. 15, 250–261. doi:10.1111/1755-0998.12302.

Sze, Y., Miranda, L. N., Sin, T. M., and Huang, D. (2018). Characterising planktonic dinoflagellate diversity in Singapore using DNA metabarcoding. Metabarcoding and Metagenomics 2, e25136. doi:10.3897/mbmg.2.25136.

Tyler, A. D., Mataseje, L., Urfano, C. J., Schmidt, L., Antonation, K. S., Mulvey, M. R., et al. (2018). Evaluation of Oxford Nanopore’s MinION Sequencing Device for Microbial Whole Genome Sequencing Applications. Sci. Rep. 8, 10931. doi:10.1038/s41598-018-29334-5.

Wick, R. R., Judd, L. M., and Holt, K. E. (2019). Performance of neural network basecalling tools for Oxford Nanopore sequencing. Genome Biol. 20, 129. doi:10.1186/s13059-019-1727-y.

Wilson, E. O. (2017). Biodiversity research requires more boots on the ground. Nature Ecology & Evolution 1, 1590–1591. doi:10.1038/s41559-017-0360-y.

Yeo, D., Srivathsan, A., and Meier, R. (2020). Longer is not always better: Optimizing barcode length for large-scale species discovery and identification. Systematic Biology. doi:10.1093/sysbio/syaa014.

Yip, Z. T., Quek, R. Z. B., Low, J. K. Y., Wilson, B., Bauman, A. G., Chou, L. M., et al. (2018). Diversity and phylogeny of Sargassum (Fucales, Phaeophyceae) in Singapore. Phytotaxa 369, 200. doi:10.11646/phytotaxa.369.3.3.

Zhang, J., Kobert, K., Flouri, T., and Stamatakis, A. (2014). PEAR: a fast and accurate Illumina Paired-End reAd mergeR. Bioinformatics 30, 614–620. doi:10.1093/bioinformatics/btt593.

