## Supplementary Table 1 and 2 for "MinION-in-ARMS: Nanopore Sequencing To Expedite Barcoding Of Specimen-Rich Macrofaunal Samples From Autonomous Reef Monitoring Structures"

Supplementary Table 1. Previous studies that deployed Autonomous Reef Monitoring Structures (ARMS) units, including the size-fraction analysed, and the method(s) used. Studies were found using search key “*Autonomous Reef Monitoring Structures*” on Google Scholar. Search results such as reviews, where ARMS were only mentioned and not deployed, were not incorporated in this table.

| **Reference** | **Size-fraction analysed** | **Method(s) used** |
| --- | --- | --- |
| Plaisance, Brainard, Caley, & Knowlton (2011a) | Motile; 1–2mm (Crustacea only) | Sanger sequencing (242 samples) |
| Plaisance, Caley, Brainard, & Knowlton (2011b) | Motile; no size fraction specified (Crustacea only) | Sanger sequencing (sample size not specified; 1,094 sequences obtained) |
| Leray and Knowlton (2015) | Motile; 106µm, 500µm and 2mm (all invertebrates)  Sessile (all invertebrates) | Sanger sequencing for 2mm samples (sample size not specified; 1,153 sequences obtained)  NGS metabarcoding for all other size fractions |
| Hurley et al. (2016) | Motile; 2mm (Brachyura only) | Morphological identification of 663 samples; no DNA sequencing was performed |
| Al-Rshaidat et al. (2016) | Motile; 106µm, 500µm and 2mm (all invertebrates)  Sessile (all invertebrates) | Sanger sequencing for 2mm sized-fraction (335 specimens; 331 sequences)  NGS metabarcoding for all other size fractions |
| Pearman, Anlauf, Irigoien, & Carvalho (2016) | Motile; 106µm, 500µm (all invertebrates); 2mm size fraction was not considered  Sessile (all invertebrates) | NGS metabarcoding |
| Pennesi and Danovaro (2017) | Sessile (Microphytobenthos) | Light microscopy and scanning electron microscopy (SEM); no DNA sequencing was performed |
| Ransome et al. (2017) | Motile; 106µm, 500µm (all invertebrates); 2mm size fraction was not considered  Sessile (all invertebrates) | NGS metabarcoding |
| Pearman et al. (2018) | Motile; 106µm, 500µm and 2mm (all invertebrates)  Sessile (all invertebrates) | Sanger sequencing (sample size and number of sequences obtained not specified; 402 OTUs reported)  NGS metabarcoding |
| Carvalho et al. (2019) | Motile; 106µm, 500µm (all invertebrates); 2mm size fraction was not considered  Sessile (all invertebrates) | NGS metabarcoding |
| David et al. (2019) | Sessile (all invertebrates) | Photograph analyses; no DNA sequencing was performed |
| Hazeri et al. (2019) | Motile; no size fraction specified (Crustacea only) | Morphological identification of 210 samples; no DNA sequencing was performed |
| Pearman et al. (2019) | Sessile (Bacteria only) | NGS metabarcoding |

Supplementary Table 2. The two demultiplexing settings used when running the unique tag mode (*-m 1*) of miniBarcoder.py.

| **Criteria** | **Full** | **Half** |
| --- | --- | --- |
| Perfect forward and reverse tag | YES | YES |
| Perfect forward, reverse mutant tag | YES | YES |
| Mutant forward, perfect reverse tag | YES | YES |
| Mutant forward and reverse tag | YES | NO |
| Perfect forward only | YES | YES |
| Perfect reverse only | YES | YES |
| Mutant forward only | YES | NO |
| Mutant reverse only | YES | NO |
